# The molecular chaperone DNAJB6 provides surveillance of FG-Nups and is required for interphase nuclear pore complex biogenesis

**DOI:** 10.1101/2021.10.26.465890

**Authors:** E. F. Elsiena Kuiper, Paola Gallardo, Tessa Bergsma, Muriel Mari, Maiara Kolbe Musskopf, Jeroen Kuipers, Ben N. G. Giepmans, Anton Steen, Liesbeth M. Veenhoff, Harm H. Kampinga, Steven Bergink

## Abstract

Biogenesis of nuclear pore complexes (NPCs) includes the formation of the permeability barrier composed of phenylalanine-glycine-rich nucleoporins (FG-Nups) that regulate the selective passage/crossing of biomolecules. The FG-Nups are intrinsically disordered and prone to liquid-liquid phase separate^1,2^ and aggregate when isolated^3^. It has remained largely unclear how FG-Nups are protected from making inappropriate interactions during NPC biogenesis. We found that DNAJB6, a molecular chaperone of the heat shock protein network, formed foci next to NPCs. The number of these foci decreases upon removal of proteins involved in the early steps of interphase NPC biogenesis. Reversely, when this process is stalled in the last steps, the number of DNAJB6-containing foci increases and they could be identified as herniations at the nuclear envelope (NE). Immunoelectron tomography showed that DNAJB6 localizes inside the lumen of the herniations arising at NPC biogenesis intermediates. Interestingly, loss of DNAJB6 results in annulate lamellae, which are structures containing partly assembled NPCs associated with disturbances in NPC biogenesis. We find that DNAJB6 binds to FG-Nups and can prevent the aggregation of the FG-region of several FG-Nups in cells and *in vitro*. Together, our data show that DNAJB6 provides quality control during NPC biogenesis and is the first molecular chaperone that is involved in the surveillance of native intrinsically disordered proteins, including FG-Nups.

---

NPCs are the sole gateways between the nucleus and the cytoplasm of the eukaryotic cell and consist of an eight-fold symmetrical cylindrical structure embedded in the double membrane of the NE. Being one of the largest protein complexes in cells, the NPC consists of over 30 different proteins, the nucleoporins (Nups) each present in different stoichiometries^4^,^5^. Assembling such a large multi-subunit structure while fusing the membranes of the NE, plus the propensity of the FG-Nups to aggregate, make NPC biogenesis a multistep and complex event. Removal of misassembled NPC intermediates or damaged NPCs is a risk to the cell as it could possibly lead to loss of compartmentalisation during disassembly^6^. In ageing, as well as in disease, it becomes clear how NPC function is intricately interwoven with cell physiology, as in both instances a loss of NPC function has been observed^6–9^. In particular, dividing cells depend on a constant supply of new NPCs, and indeed NPC assembly is compromised in mitotically aged yeast cells^8^.

During our studies on the molecular chaperone DNAJB6 we noticed that DNAJB6 localizes to specific foci at the NE (Fig. 1A and S1A). DNAJB6-containing foci colocalise with LaminB1 (Fig. 1B) and are in close proximity to NPCs (Fig. 1C). To validate our antibody we stained DNAJB6 knockout (KO) cells and the foci at the nuclear rim were absent (Fig. 1D). Strikingly, DNAJB6 KO cells contain large accumulations of NPC material in the cytoplasm (Fig. 1D, 1E), whereas LaminB staining is unaffected, suggesting that the NE is still intact (Fig. S1B). In addition, NPC components in wild type cells are mostly homogeneously distributed throughout the NE, whereas they are irregularly distributed over the NE in DNAJB6 KO cells (Fig. S1C). Knockdown of DNAJB6 using siRNA results in similar accumulations of NPC-positive material in the cytoplasm (Fig. S1D), while adding back GFP-DNAJB6b reversed this (Fig. S1E). Transmission electron microscopy (TEM) in DNAJB6 KO cells revealed that the cytoplasmic accumulations are actually NPC-like structures aligned in sheets (Fig. 1F) or in strings (Fig. S1F), resembling annulate lamellae^10,11^. The annulate lamellae that accumulate in DNAJB6 KO cells lack nucleoporins from the basket, but stain positive for components of other NPC subcomplexes (Fig. 1G and 1H), which is similar to annulate lamellae described by others^12–16^. The appearance of annulate lamellae in somatic cycling cells has been associated with disruptions in NPC biogenesis^16,17^ that can lead to disturbances in nucleocytoplasmic shuttling, including the mislocalization of RNA^7,18,19^. Indeed, DNAJB6 KO cells show a retention of RNA in the nucleoplasm (Fig. S1G).

**Figure 1.**
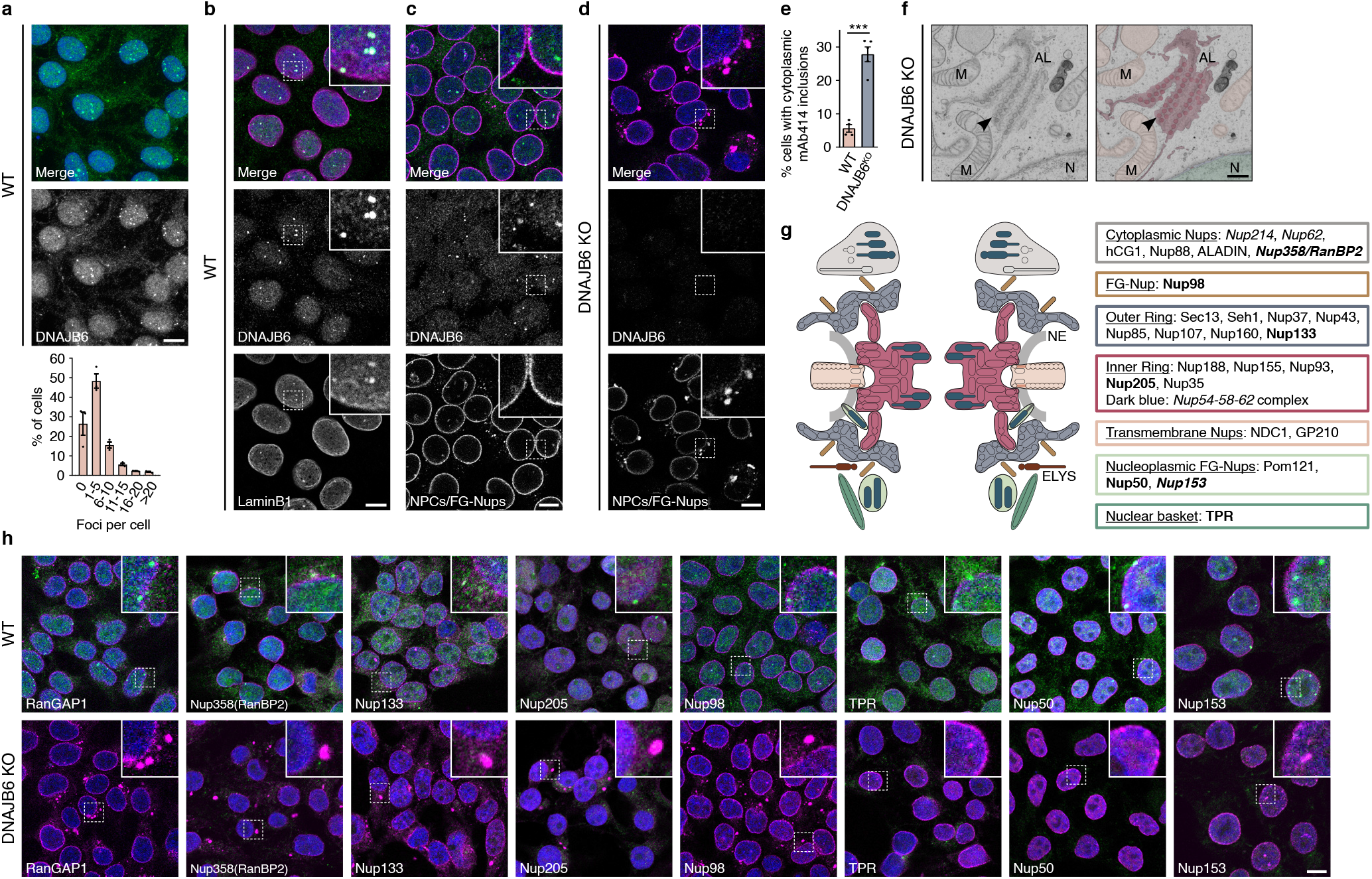
DNAJB6 localizes in foci to the nuclear envelope and disruption of DNAJB6 expression leads to accumulation of nuclear pore complexes (NPCs) in the cytoplasm. **A.** Z-projection of 30 consecutive slices of 0.5μm of HEK293T cells stained for DNAJB6. Cells show nuclear and cytoplasmic staining of DNAJB6, as well as brighter foci located at the nuclear envelope. Graph depicts a quantification of number of DNAJB6 foci per cell; mean±SEM. In HEK293T cells over 95% of cells have between 0 and 15 foci at the NE at a given time. **B.** Z-projection of 20 consecutive slices of 0.5μm of HEK293T cells stained for DNAJB6 (green) and LaminB1 (magenta) shows colocalization of LaminB1 with DNAJB6 foci. **C.** HEK293T WT cells stained for DNAJB6 (green) and NPCs/FG-Nups with mAb414 (magenta). DNAJB6 foci localize in close adjacency to NPCs at the nuclear envelope (NE) and show normal localization of NPC components to the NE. **D.** HEK293T DNAJB6 KO cells stained for DNAJB6 (green) and NPCs (mAb414, magenta) have distinct cytoplasmic accumulations of NPCs. **E.** Quantification of percentage of cells with cytoplasmic NPC accumulations ***p≤0.001; n≥4 independent samples, unpaired t-test, mean±SEM. **F.** Cytoplasmic NPC accumulations or annulate lamellae (AL) visualised by transmission electron microscopy and AL-NPCs pseudo-colored in bright pink, can be distinguished in cytoplasmic sheets. Arrows point at individually identifiable NPCs. Nucleus indicated with N, mitochondrion indicated with M. Scale bar represents 500nm. **G.** Schematic overview of the composition of the NPC, adapted from: Fernandez-Martinez & Rout, Trends Biochem. Sci. 2021. 46(7):595-607. Proteins with FG-repeat domains are dark blue and Nup98 in light brown. Indicated in italic are NPC components stained by the mAb414 antibody. Indicated in bold are NPC components stained for in panel H. **H.** Associated proteins on the cytoplasmic side of the NPC (RanGap1), and different nucleoporins of the NPC of cytoplasmic filaments (Nup358), the outer ring (Nup133), the inner ring (Nup205), FG-Nups (Nup98), and the intranuclear basket (TPR, Nup50, Nup153) are stained (magenta) with DNAJB6 (green) in WT and DNAJB6 KO cells. Scale bars on all fluorescent images represent 10μm.

DNAJB6 belongs to the large class of J domain proteins (JDPs) that function as co-chaperones of the HSP70 chaperones. To see if other JDPs are also present at the NE and lead to annulate lamellae formation, we tested a subset of them using ectopically expressed GFP-tagged versions. The closely related DNAJB2 and testis-specific DNAJB8^20^ are also found to localize at the NE where they colocalize with DNAJB6 foci (Fig. S1H). The more dissimilar JDPs, GFP-DNAJB1 and GFP-DNAJA1^21^, do not (Fig. S1H). Moreover, knockdown of DNAJB2 also results in the formation of annulate lamellae, whereas knockdown of DNAJB1 and DNAJA1 do not (Fig. S1I). This argues that formation of annulate lamellae is specific for the absence of a subclass of JDPs to which DNAJB6 belongs.

In higher eukaryotes, which have open mitosis, there are two main mechanisms of NPC biogenesis: 1) post-mitotic biogenesis when the NE and NPCs in both nuclei reassemble after cell division, and 2) interphase biogenesis when new NPCs are inserted into the double membrane of the NE^22^. By synchronizing cells in G1/early S-phase using a double thymidine block (Fig. 2A top and S2A), we noted that DNAJB6 foci decrease at the start of mitosis, and foci numbers only increase when cells have progressed through mitosis and enter G1/S (Fig. 2A bottom and S2B). Thus, DNAJB6 foci correlate with interphase biogenesis as they appear in G1/S when nuclear envelopes are re-established and post-mitotic NPC biogenesis in telophase has already been completed. To corroborate this, we knocked down the critical nucleoporins for either post-mitotic or interphase assembly. Knockdown of ELYS, required for the initiation of post-mitotic NPC biogenesis^23^, did not result in changes in the number of DNAJB6 foci (Fig. 2B panel 2). In contrast, knockdown of Nup153, the basket FG-Nup that is an important player in initiation of interphase NPC biogenesis^24^, results in a significant drop in the number of DNAJB6 foci at the NE (Fig. 2B panel 3). Knockdown of ELYS as well as Nup153 also induce annulate lamellae, while knockdown of the cytoplasmic NPC component Nup358 did not (Fig. S2C). This is in line with both ELYS and Nup153 being needed for NPC biogenesis, albeit in a different step of the cell cycle^23,24^.

**Figure 2.**
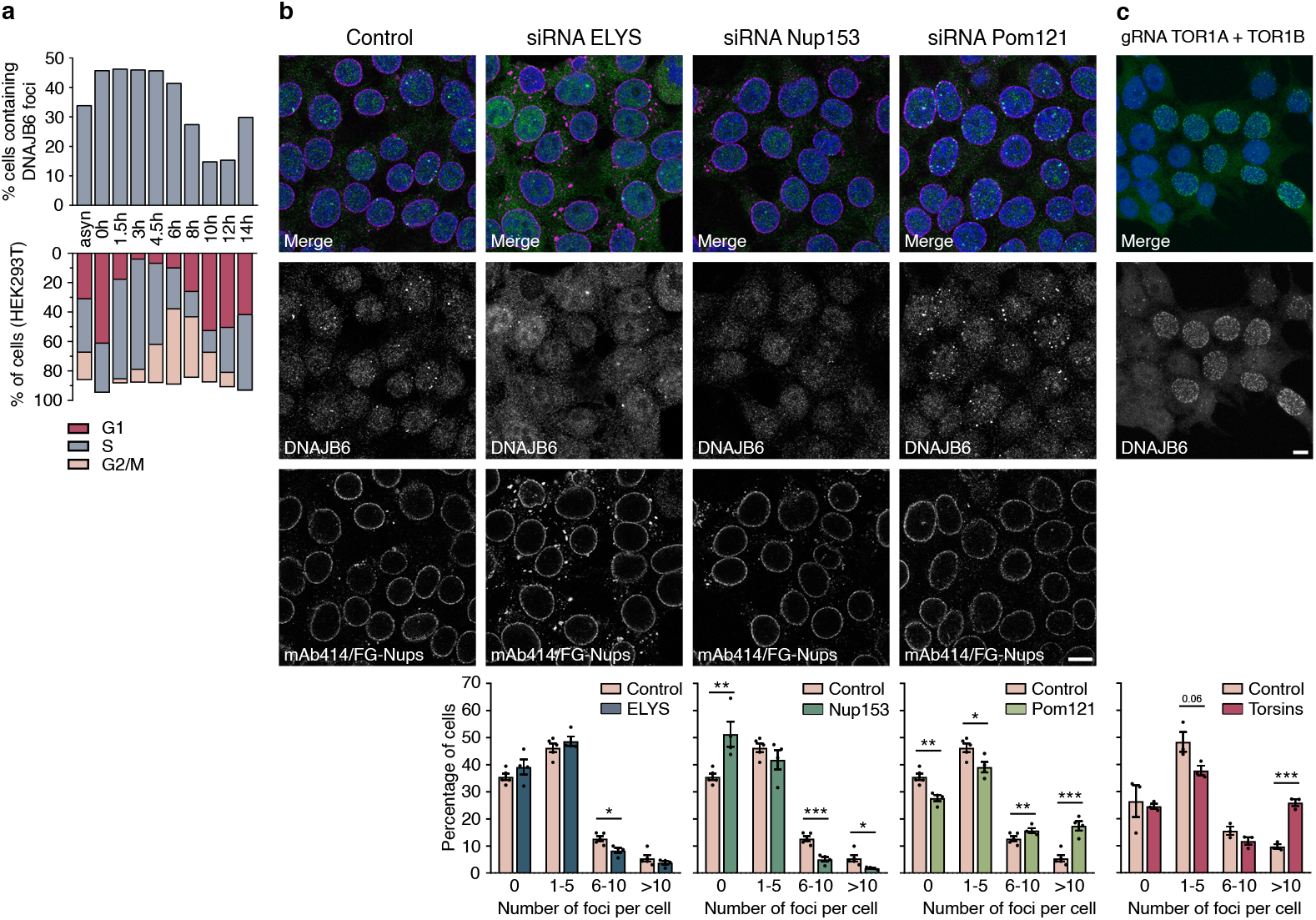
DNAJB6 foci formation is cell cycle dependent and is related to interphase NPC assembly. **A.** Top panel depicts percentage of DNAJB6 foci in HEK293T cells after release from cell cycle synchronization. In asynchronous cells (asyn.) ~40% of cells contain DNAJB6 foci. The percentage of foci-containing cells is given for the depicted time after the release from the double thymidine block. Lower panel shows synchronization profile of HEK2393T cells after being synchronized in G1/S-phase of the cell cycle by a double thymidine block. **B.** DNAJB6 foci (green) at the NE (NPC staining mAb414, magenta) upon siRNA mediated knockdown of the nucleoporin ELYS, Nup153, or Pom121. Graphs depict a distribution of the amount of DNAJB6 foci per cell for the indicated siRNA treatment. NB: these conditions all have the same control. *p≤0.05, **p≤0.01, ***p≤0.001; n≥4 independent samples with total >5500 cells counted per condition, automated with ICY; mean±SEM. **C.** Z-projection of 30 consecutive slices of 0.5μm of HEK293T cells stained for DNAJB6 (green) with CRISPRi knockdown of Torsin A and B, with a distribution of amount of DNAJB6 foci per cell. Upon knockdown of Torsin A and B, cells with higher amount of foci increases to ~25%. ***p≤0.001; n=3 independent samples with total >450 cells counted, automated with ICY; mean±SEM. Scale bars on all fluorescent images represent 10μm.

To more precisely establish in which step of NPC biogenesis DNAJB6 foci appear, we knocked down Pom121 which blocks the process of fusion of the NE^25–27^, leading to a block in late interphase NPC biogenesis. In Pom121 depleted cells, we observe a significant increase in the number of DNAJB6 foci (Fig. 2B panel 4), indicating that DNAJB6 is involved in a step in NPC biogenesis prior to the fusion of the NE. To further test if DNAJB6 foci are associated with (stalled) NPC intermediates, cells were depleted of TorsinA and TorsinB with CRISPRi (Fig. 2C). Torsins are thought to be responsible for completing interphase NPC biogenesis, possibly by playing a role in inner nuclear membrane (INM)/outer nuclear membrane (ONM) fusion^28^. In line with Pom121 depletion, depletion of Torsins also resulted in an increase in the number of DNAJB6 foci per cell (Fig. 2C and Fig S2D). The functionality of Torsins depends on their ability to form heterohexameric complexes with one of the co-factors LAP1 or LULL1^29,30^. Interestingly, ectopic expression of a mutant TorsinA that cannot interact with its co-factors (TorsinΔE302^30^), and is causative for early-onset DYT1 dystonia^31^, results in an increase in DNAJB6 foci as well (Fig. S2E). Moreover, Torsin-induced foci at the NE are known to be ubiquitin positive^29^ and we confirm the presence of ubiquitin in the DNAJB6 foci (Fig. S2F). Together, these results show that DNAJB6 foci coincide with interphase NPC biogenesis and point to a role of DNAJB6 at an early stage of the process, i.e. after initiation (Nup153) but prior to fusion of the pre-assembled cytoplasmic subcomplexes with the NE.

Stalling NPC biogenesis in interphase by knocking down Torsins leads to the formation of herniations at the NE^32^ (Fig. 3A). Transmission electron microscopy revealed that both endogenous DNAJB6 (Fig. 3A, left) and ectopically expressed GFP-DNAJB6b (Fig. 3A right) localize within these herniations. At the neck of the herniations, a structure reminiscent of the NPC is recurrently observed^28,33^. FG-Nups, labelled with mAb414 antibody, localize to NPCs that are embedded in the NE (Fig. 3B left) and in rare occasions can be found at the neck of a herniation (Fig. 3B right). Double labelling for mAb414 (10nm gold) and GFP (15nm gold) confirms the localization of NPC components at the NE and GFP-DNAJB6b within the herniations (Fig. 3C). Herniations have also been suggested to carry specific cargo^34,35^, and in a recent study it was indeed shown that the content is not random, as no canonical ERAD^33^ and no nuclear transport or ribosome components^28^ were found. Importantly, DNAJB6, DNAJB2, and other HSP70 machinery proteins, were identified in a mass spectrometry analysis of nuclear envelopes isolated from Torsin depleted cells^28^. In that study^28^, Myeloid Leukemia Factor 2 (MLF2) was also identified as a major component of herniations. Consistently, we find both HSP70 (HSPA1A) and MLF2 to colocalize with DNAJB6 foci at the NE (Fig. S3A and S3B).

**Figure 3.**
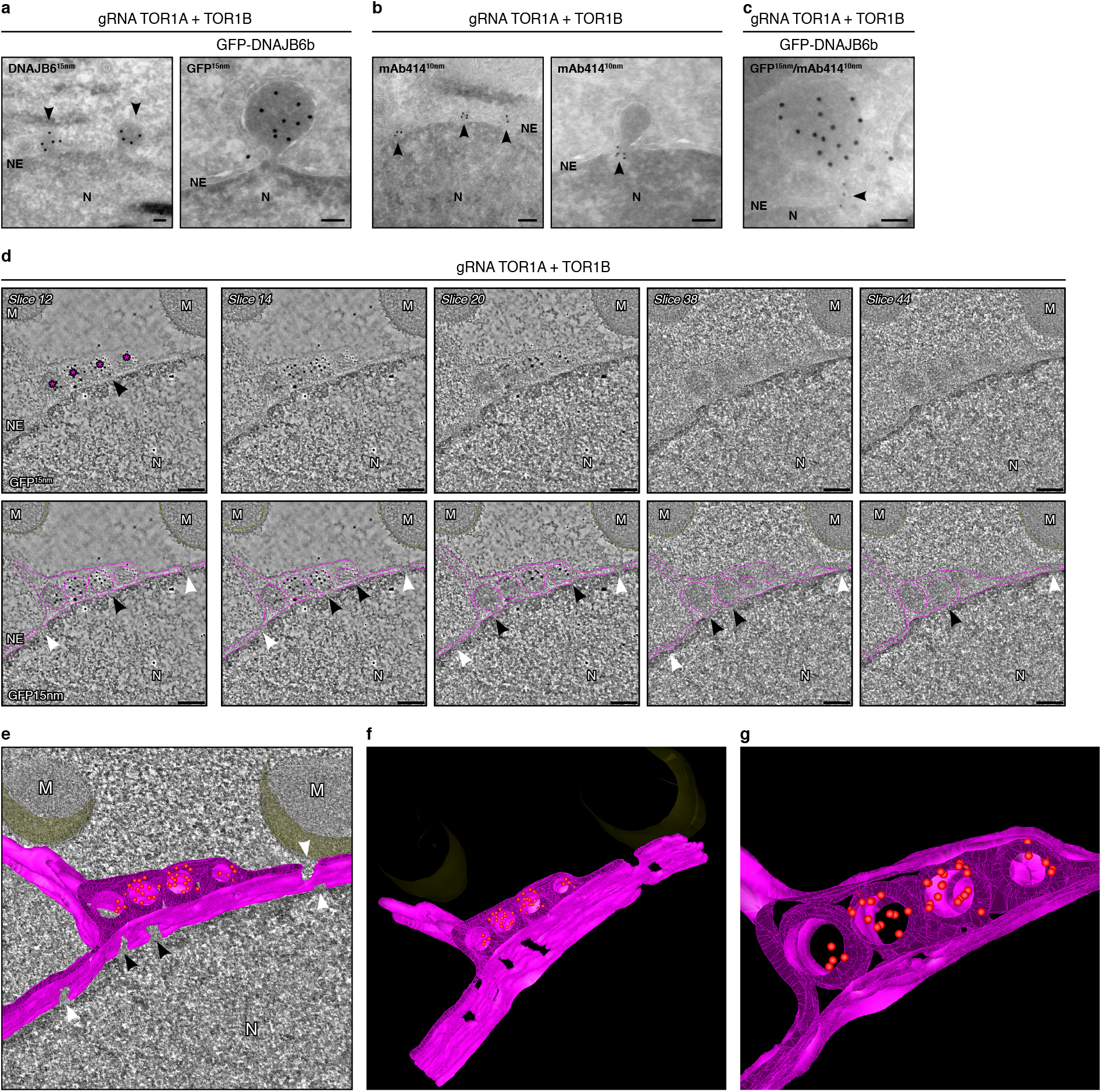
DNAJB6 localizes to herniations at the nuclear envelope. **A.** Transmission electron microscopy images of HEK293T cells with CRISPRi knockdown of Torsin A and B, either without expression of GFP-DNAJB6b (left, arrowheads indicate herniations) or with GFP-DNAJB6b (right), processed with Tokuyasu cryosectioning. Cryosections of 250 nm thickness were labelled with either the anti-DNAJB6 or anti-GFP antibody, followed by incubation with 15 nm gold particle-conjugated protein. **B.** Cryosections were labelled with mAb414 antibody against FG-Nups, followed by incubation with 10 nm gold particle-conjugated protein A, showing clear labelling of NPCs (left). In rare cases labelling can be observed at the neck of herniations (right). **C.** Double labelling of sections for GFP-DNAJB6b and FG-Nups shows proximity of DNAJB6 in a herniation to the NPC. All scale bars in panels A, B and C represent 500nm. Nucleus is indicated with N, nuclear envelope indicated with NE. **D.** On immunoelectron to-mography slices surface labelling was used to select areas of interest prior to recording tilt series. Tomographic slice (Z slice 12) revealing the specific immunogold (15 nm) labelling on the surface of the cryosection. Asterisk indicating herniations. Indicated tomographic slices extracted from the tomogram showing herniations at the NE. In the bottom images, nuclear membranes are traced (pink). More electron dense areas can be observed at the neck of the herniations where the NE bends into the herniation, indicated by black arrow heads. NPCs are indicated with white arrow heads. Nucleus is indicated with N, mitochondria with M, and nuclear envelope with NE. Scalebars represent 156nm. **E.** Three-dimensional model of GFP-DNAJB6b-positive herniations were obtained by electron tomography of 400 to 450 nm thick cryosections. The NE is represented in pink, and GFP gold labelling in red. Openings underneath herniations are indicated with black arrow heads, and NPCs with white arrow heads. **F. and G.** Stills from the 3D model showing in **F.** a view from the nuclear side where the openings of the NPCs and the herniations can be seen and in **G.** a close-up of the herniations with the localization of the GFP gold labelling in red.

Next, we used immunoelectron tomography on Tokuyasu thick cryosections^36^ to obtain insight in the 3D organisation of the DNAJB6-containing herniations. The herniations (Fig. 3D, indicated with asterisks on slice 12) were located in the cell sections by immuno-gold labelling using GFP-antibody (Fig. 3D) or an antibody against DNAJB6 (Fig. S3C). Subsequent z-slices obtained through the tomography of the GFP-DNAJB6b-positive herniations reveal that the double membrane of the NE continues into the membrane around the herniation (Fig. 3D, 3E, and Video 1). On some occasions, several individual INM herniations are enclosed by a single ONM. The INM surrounding these individual herniations were never found to be fused with INMs of other herniations (Fig. 3G). Recently, herniations with NPC-resembling structures at the cytoplasmic side have been described to occur under starvation conditions, and were proposed to involve NPC turnover^37^. From the tomography it becomes evident that the ONM of the herniations does not have openings and is continuous with the ER (Fig. 3D). In addition, starvation did not induce DNAJB6 foci formation (data not shown). These two findings suggest that the DNAJB6-containing herniations are not related to (starvation-induced) NPC turnover. Our data is in line with a role of DNAJB6 in interphase NPC biogenesis since the 3D model of the DNAJB6-containing herniations clearly shows that there are only openings underneath the herniations at the NE (Fig. 3D, black arrows). At the neck, it contains a more electron dense structure with similar dimensions as NPCs in the NE (Fig. 3E and 3F). Together with the NPC labelling (Fig. 3B and 3C), this suggests that NPC material is present at the neck of the herniations and is in line with the presence of an NPC biogenesis intermediate^33^.

Treating the cells with the aliphatic alcohol 1,6-hexanediol, which is known to disrupt the permeability barrier of the NPC *in vivo*^38^ and to disrupt liquid-liquid phase interactions of FG-Nups *in vitro*^39^, almost immediately leads to a loss of DNAJB6 foci at the NE (Fig. S3D). DNAJB6 foci largely remained after treatment with 2,5-hexanediol, which has little effect on the melting of FG domain polymers (Fig. S3D). We speculate that when INM/ONM fusion is stalled, herniations are formed that contain DNAJB6, and possibly other factors involved in NPC biogenesis. These factors would normally have been released into the cytoplasm if NPC biogenesis and fusion of INM/ONM would proceed as usual, but now they are caught inside the herniations. The presence of 1,6-hexanediol allows DNAJB6 to exit the herniations on the nuclear side resolving the foci. These data are consistent with an inside-out extrusion of the INM from the nucleus towards the ONM, preceding the completion and insertion of the NPC into the NE as has been suggested before^40^.

DNAJB6 has been shown to prevent aggregation of amyloidogenic substrates including polyglutamine (polyQ)-containing proteins^41–43^, amyloid-β^44^, α-synuclein^45^, and yeast prions^46,47^. The early presence of DNAJB6 at sites for NPC biogenesis could reflect the ability of this chaperone to specifically prevent FG-Nups to undergo unwanted interactions, and possibly even aggregate^3^, during biogenesis. By using a BioID2-DNAJB6b fusion construct, several FG-Nups and importin β could be identified as close proximity partners of DNAJB6b (Fig. 4A, S4A, and S4B), but not Nup133, a component of the outer ring complex and not an FG-Nup (Fig. 4A). One of the FG-Nups that we identified by using the mAb414 antibody is Nup153 (Fig. 4A), a key FG-Nup in initiating interphase biogenesis^24^, and implicated in DNAJB6 foci formation (Fig. 2B). To test if Nup153 may be a client of DNAJB6, we overexpressed the FG-region of human Nup153 (hNup153FG) tagged with GFP, which can undergo the transition into an amyloidogenic state *in vitro*^3^. Two categories of accumulations are observed: smaller spherical shaped and larger fibrous structures (Fig. 4B). The ability to form such accumulations is specifically related to the presence of the FG repeats as GFP-Nup153AG - a mutant in which all phenylalanines have been replaced with alanines - does not appear to form accumulations of any kind^3^ (Fig. S4C). DNAJB6 colocalizes with hNup153FG (Fig. S4D) and, more importantly, the percentage of cells with large fibrous structures increase in a DNAJB6 KO background, while the formation of fibrous structures is almost completely suppressed upon DNAJB6b overexpression or re-expression (Fig. 4C). The different accumulations found for hNup153FG have distinct properties. The spherical shaped accumulations can be characterized as mobile as they show quick fluorescence recovery after photobleaching (FRAP) (Fig. 4D and 4E top). Within these accumulations, fluorescence is quickly redistributed after bleaching and fluorescence is even regained almost back to pre-bleached levels, suggesting exchange with its surroundings. However, the larger fibrous structures do not recover from photobleaching and have a more solid character (Fig. 4D and 4E bottom). In addition, a small fraction of hNup153FG is 0.5% SDS insoluble and is retained on 0.2μm pore size cellulose acetate filters, indicative of amyloid formation (Fig 4F and 4G). Similar to what we find for the larger fibrous structures, the amount of SDS-insoluble material of hNup153FG increases in DNAJB6 KO cells and is reduced upon DNAJB6 overexpression (Fig. 4F and 4G). Amyloid formation of hNup153FG strictly depends on the FG-motifs, as the GFP-Nup153AG mutant does not form any visible condensates or aggregates (Fig. S4D) and is not retained on the cellulose acetate filters after SDS extraction (Fig. S4E), as was shown before^3^. Moreover, the association between hNup153FG and DNAJB6 (Fig. S4F, last lane) also specifically depends on the presence of the FG-motifs (Fig. S4F, middle lane).

**Figure 4.**
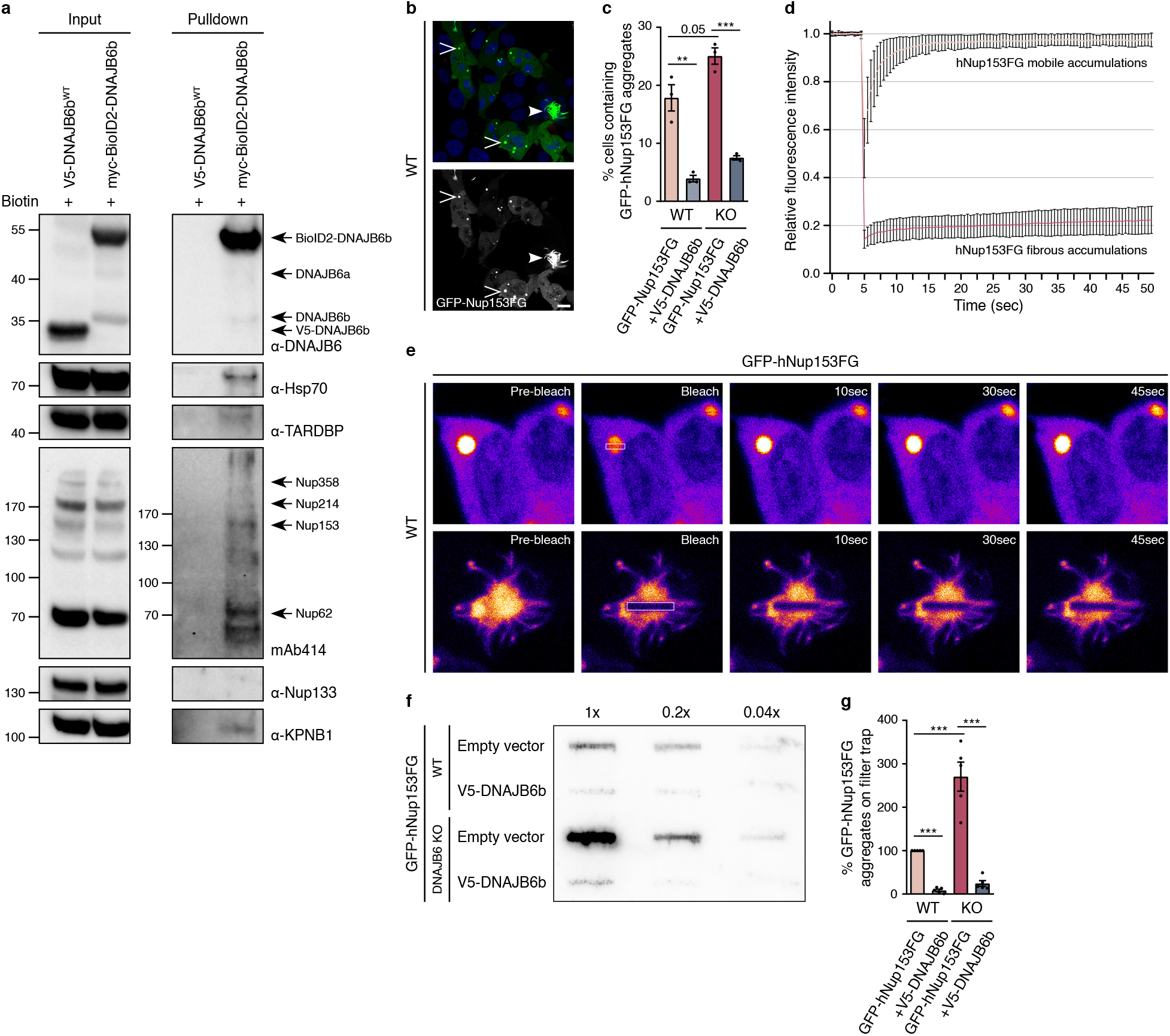
DNAJB6 interacts with FG-Nup components of the nuclear pore complex. **A.** HEK293T cells expressing V5-DNA-JB6b or BioID2-DNAJB6b, treated for 24h with 50μM biotin. Biotinylated close proximity partners of DNAJB6b were pulled down by biotin affinity immunoprecipitation with streptavidin beads, run on SDS-PAGE, and nucleoporins were detected with specific antibodies against Nup133 and mAb414. **B.** HEK293T expressing GFP-hNup153FG show spherical accumulations (open arrow-head) and fibrous aggregates (closed arrowhead). Scale bar represents 10μm **C.** Percentage of cells containing fibrous structures of GFP-hNup153FG in WT and DNAJB6 KO cells without and with co-expression of V5-DNAJB6b. **p≤0.01, ***p≤0.001; n=3 independent samples; mean±SEM. **D.** Fluorescence Recovery After Photobleaching (FRAP) on spherical GFP-hNup153 FG accumulations shows quick recovery after photobleaching (mobile accumulations), whereas fluorescence of fibrous structures does not recover fluorescence in bleached areas. FRAP curves with error bars representing SD over three independent experi-ments with N=35 for mobile and N=14 for fibrous structures. **E.** Fluorescence images of FRAP on spherical (mobile) and fibrous (immobile) structures formed by GFP-Nup153FG. **F.** Filter-trap assay of cells with GFP-hNup153FG overexpression. Three serial 5-fold dilutions were loaded onto cellulose-acetate membranes and probed with anti-GFP antibodies to detect aggregation of the GFP-hNup153FG fragment in either HEK293T WT or DNAJB6 KO cells and without or with co-overexpression of V5-DNAJB6b. **G.** Band intensities of the data in **F** are quantified relative to the intensity of aggregation of hNup153FG in WT HEK293T. ***p≤0.001; n=5 independent samples; mean±SEM.

To test if DNAJB6 is directly responsible for the anti-aggregation effects on hNup153FG, we purified DNAJB6b and the FG-region of hNup153. Recombinant hNup153FG becomes insoluble, depending on the crowding reagent used, and can be trapped on a filter trap, indicating it is aggregated (Fig. S5). Importantly, adding recombinant DNAJB6b reduces the aggregation of hNup153FG in a dose dependent manner (Fig. 5A). We also tested the activity of DNAJB6b on the aggregation of the FG-regions of the yeast Nup100, Nup116, and Nup145N. yNup100FG, yNup116FG, and yNup145NFG form aggregates as well, and this aggregation is strongly reduced in the presence of DNAJB6b (Fig. 5A). The anti-aggregation activity of DNAJB6b is specific, as neither BSA, MBP-mCherry, nor DNAJA1 influence the aggregation of either hNup153FG or yNup100FG (Fig. 5B). DNAJB6b delays the aggregation kinetics (Fig. 5C and Fig. 5D). In the presence of DNAJB6b, the aggregation of both hNup153FG and yNup100FG are only slightly increasing at 120 minutes, whereas aggregation in the absence of DNAJB6b is already saturated at that time point. These results establish the direct anti-aggregation effect of DNAJB6b on the FG-region of different FG-Nups.

**Figure 5.**
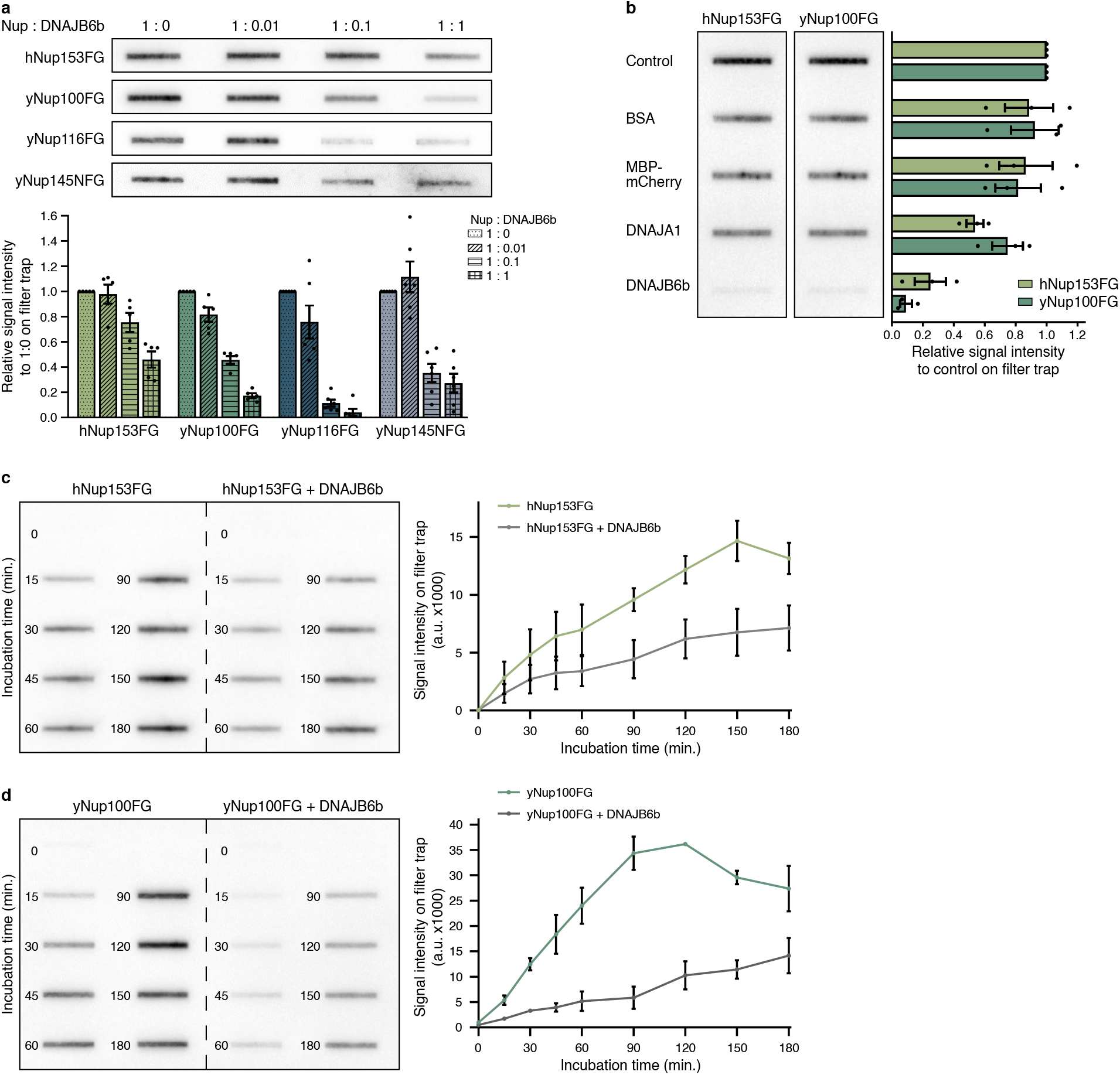
DNAJB6b prevents the aggregation of human Nup153FG and yeast Nup100FG, Nup116FG and Nup145NFG *in vitro*. **A.** Filter trap assay to detect aggregation of 3 μM of purified FG-Nup fragments, incubated with indicated molar ratios of DNAJB6b, probed with anti-His antibodies to detect FG-Nup fragments. hNup153FG, yNup100FG, and yNup116FG were incubated in the assay buffer (150mM NaCl, 50mM TrisHCl, pH8) containing 10% PEG3350 (w/v), at 25°C during either 1 hour (hNup153FG, yNup100FG) or 3 hours (yNup116FG), and yeast Nup145NFG was incubated in a buffer comprising 50mM NaCl, 100mM sodium phosphate, pH6, 10% PEG3350 (w/v), at 25°C during 1 hour. Band intensities on filter trap are quantified relative to the intensity in the absence of DNAJB6b. n=5-6 independent samples; mean±SEM. **B.** As in A, 3 μM of purified hNup153FG or yNup100FG were incubated in the absence (control) or presence of BSA, MBP-mCherry, DNAJA1, or DNAJB6b at a 3 μM concentration (molar ratio 1:1). BSA and MBP-mCherry served as non-chaperone controls and DNAJA1 served as chaperone control. Band intensities are quantified relative to the intensity of the FG-Nup fragments alone (control). n=3 independent samples; mean±SEM. **C, D.** Conditions as in A, showing here the time-dependent formation of hNup153FG (**C**) and yNup100FG (**D**) aggregates in the absence and presence of DNAJB6b at a molar ratio of 1:1. Time is indicated in minutes. Quantification of filter trap band intensity (in arbitrary units, a.u.). n=3 independent samples; mean±SEM.

Altogether, our data show that the molecular chaperone DNAJB6 plays a prominent role in interphase NPC biogenesis and it is the first time that a chaperone of the heat shock protein network is now implicated in this process. Compromising DNAJB6 leads to disturbed NPC biogenesis, with annulate lamellae formation and problems in RNA transport. When interphase NPC biogenesis process is stalled, DNAJB6 (and other components of the HSP70 chaperone network) ends up in the resulting herniation at the NE. It is likely that if NPC biogenesis proceeds normally, the early interactors, including DNAJB6, of these NPC assembly intermediates are released at the cytoplasmic side of the NPC. Similar NE herniations have been found in DYT1 dystonia patients. We show that DNAJB6 protects the natively disordered FG-region of FG-Nups from transitioning into an amyloidogenic state. Intriguingly, in a parallel paper by the Schlieker group, the herniation co-resident MLF2 was also identified to act on the FG-Nups by modulating their interactions. This suggests that surveillance of intrinsically disordered polypeptides may be important in DYT1 disease etiology. The activity of DNAJB6 towards native FG-Nups is similar to its previously described unique anti-aggregation activity on mutant disease-related protein substrates, including polyQ-containing proteins^41–43^, and α-synuclein^45^, of which some have been suggested to be disordered as well. This makes DNAJB6 the first chaperone for natively disordered proteins in normal cellular physiology, as well as in disease. Many of the DNAJB6 substrates relate to neurodegenerative diseases^41–43,47,48^ and several are also characterized by disturbances in NPCs and/or nucleocytoplasmic transport^7,49–54^, similar to what we find as a result of DNAJB6 depletion. This suggests an intriguing link between the HSP chaperone network, ageing-related aggregation-prone proteins, and NPC assembly and function.

## Acknowledgments

EFEK is supported by a topmaster fellowship from the Groningen University Institute for Drug Exploration (GUIDE). HHK and SB are supported by a grant from the CTH (grant no. 686728). LMV, HHK, SB, MKM, PG, TB, and AS are supported by an NWO Groot (grant no. 685709). LMV, PG, TB, and AS are supported by a Vici grant (VI.C.192.031). MM is supported by an ALW Open Programme (ALWOP.355). We want to thank Jeanette Brunsting and Amarins Blaauwbroek for practical assistance. We want to thank Edward Lemke for providing us with the GFP-Nup153AG construct, Dirk Görlich for the yNup116FG and yNup145NFG constructs, and Christian Schlieker for the HeLa 4TorKO cell line. Fluorescence and live cell imaging was performed in the UMCG Microscopy and Imaging Center (UMIC), sponsored by ZonMW grant 91111.006 and NWO 175-010-2009-023. Electron microscopy 2D-imaging was performed at the UMCG microscopy and imaging center (UMIC). Tomography imaging was performed at the EM facility of the faculty of Science and Engineering of Groningen, the Netherlands.

## Author Contributions

E.F.E.K., H.H.K., L.M.V., and S.B. conceived the project. All the experiments were designed, performed, and analysed by E.F.E.K., except for the transmission electron microscopy in Fig. 1 which was performed by J.K., the electron microscopy and tomography in Fig. 3, which was performed by M.M., and the *in vitro* data in Fig. 5, which was performed by P.G.P., T.B., and A.S. Supervision was done by H.H.K., L.M.V., A.S., B.N.G., and S.B. Experimental and analysis assistance was provided by M.K.M. The manuscript was written by E.F.E.K and S.B. with input of all authors.

## Materials and methods

### Cell lines, cell culture, and transient transfections

HEK293T, DNAJB6 CRISPR-Cas9 knockout HEK293T, U2OS, HeLa, and HeLa 4TorKO cells were cultured according to standard protocols in in DMEM (GIBCO) supplemented with 10% FCS (Greiner Bio-one), and penicillin/streptomycin (Invitrogen). BeWo cells were grown in F12 (GIBCO) plus 10% FCS (Greiner Bio-one), penicillin/streptomycin (Invitrogen). For transient transfections, cells were grown to 70%–80% confluence in a 37°C incubator at 5% CO2, in 35 mm diameter dishes coated with 0.0001% poly-L-lysine (Sigma) and/or on coated coverslips for confocal microscopy analysis. For live cell imagining, cells were grown in 35 mm diameter glass-bottom wells (MatTek). For siRNA, CRISPRi, and cDNA plasmids, cells were transfected with Lipofectamine2000 (Invitrogen) according to the manufacturer’s instructions.

### Gene cloning and generation of mutants

Information about the (construction of) the V5-DNAJB6 wildtype is described in Hageman et al. 2010^41^ and Kakkar et al. 2016^42^. GFP-DNAJB6b was made through PCR of GFP with HindIII and BamHI restriction sites to replace the V5-tag of V5-DNAJB6b. To make the BioID2-tagged DNAJB6b, DNAJB6b was amplified from the V5-DNAJB6b vector and digested with NotI and BamHI and inserted into myc-BioID2 pcDNA3.1 (74223; Addgene). TorsinA-GFP (32119) was obtained from Addgene and cloned into the mammalian expression pcDNA3.1 vector with restriction sites HindIII and BamHI. The ER localization signal was reinserted using oligos. The TorsinA ΔE mutation was made by site directed mutagenesis (Agilent, 200523). MLF2-GFP was created by PCR of MLF2 from cDNA extracted from HEK293T cells and inserted into the pEGFP vector. GFP-Nup153 (87340) was obtained from Addgene. GFP-Nup153FG (aa 875-1475) was created by PCR of the FG-domain of Nup153 and inserted into the pEGFP vector. The GFP-Nup153AG construct was kindly provided by Edward Lemke.

For protein purification, the cDNA fragment encoding human Nup153FG (hNup153 FG), and yeast Nup100 (yNup100FG, aa 1-580), and the full-length chaperones DNAJB6b and DNAJA1 were cloned in the expression plasmid pSF350, containing a His6-tag at the N-terminal end and a cysteine residue at the C-terminal end. The constructs expressing yeast Nup116 (yNup116FG, aa 1-725) and yeast Nup145N (yNup145N FG, aa 1-219) were kindly provided by Dirk Görlich.

### CRISPRi

CRISPRi was performed following the protocol from Matthew H Larson, 2013^55^. Briefly, dCas9-KRAB (71237) and sgRNA (44248) were obtained from Addgene. The dCas9-KRAB was cloned into the mammalian expression vector pcDNA 3.1. The mCherry tag in the sgRNA vector was replaced by an mTurquoise to make full use of the color spectrum for immunofluorescence. sgRNA was designed by CRISPR-ERA v1.2 (http://crispr-era.stanford.edu/) following the standard protocol. sgRNAs sequences for the Torsins are: TorsinA: CGGCGCGAGAACAAGCAGGG, TorsinB: CTTCGAGGAGCGGGATGTTG. HEK293T cells in 6well plates were transfected using Lipofectamine2000 according to the manufacturer’s protocols with 1μg Cas9-KRAB plasmid and total of 1.5 μg sgRNA plasmid(s). Cells were either fixed for immunofluorescence or electron microscopy.

### BioID2 pulldown

BioID2-DNAJB6b transfected HEK293T cells were incubated for 24 hours in complete media supplemented with 50μM biotin. After two ice cold PBS washes, cells were scraped in 1ml PBS and centrifuged for 5 minutes at 1000rpm at 4°C. The PBS was removed and pellets were snap frozen in liquid nitrogen. After thawing on ice, cells were lysed in 250μl lysis buffer (150mM NaCl, 50mM Tris pH 8.0, 0.2% NP-40, 0.2% SDS, 1x Complete protease inhibitor EDTA free (Roche), and ~30 units Benzonase (EMD Millipore)), vortexed and put on ice. Lysates were vortexed every 10 minutes for an hour to assure full lysis. Lysates were centrifuged for 10 minutes at 5000rpm at 4°C and supernatants were transferred to new tubes and pellets discarded. Protein concentrations were equalized in washing buffer (150 mM NaCl, 50 mM Tris pH 8.0, 1mM EDTA, 0.2% NP-40, 0.2% SDS, 1x Complete protease inhibitor (Roche)) and 50μl put in new tubes labelled ‘input’ with 25μl of 4x Laemli buffer with 20% β-mercaptoethanol and immediately boiled for 5 minutes. To the rest of the samples 30μl of prewashed (2x in 500μl washing buffer) magnetic streptavidin beads (Invitrogen) were added and incubated on a roller in a 50ml tube at 4°C overnight. All subsequent steps at room temperature. Supernatant was removed (50μl was taken for the ‘supernatant’ fraction) and beads were washed twice in 500μl washing buffer, twice in 500μl high salt buffer (500 mM NaCl, 50 mM Tris pH 8.0, 1mM EDTA, 0.2% NP-40, 0.2% SDS), and twice in 500μl washing buffer. 50μl of 4x Laemli buffer with 20% β-mercaptoethanol was added and beads were incubated for 15 minutes at 70°C. Sample was removed from beads and boiled for 5 minutes, labelled ‘pulldown’. 5μl input and 10μl pulldown were loaded for western blotting and probed with indicated antibodies.

### Immunofluorescence

For immunofluorescence, cells were washed once with PBS to remove residual media and fixed using 2% paraformaldehyde (PFA) in PBS for 15 minutes. After washing twice with PBS, cells were permeabilized with 0.1% Triton X-100 for 5 minutes at room temperature and washed again with PBS. Then the cells were washed twice in PBS+ (PBS complemented with 0.5% bovine serum albumin and 0.15% glycine) and incubated with 35μl primary antibody in PBS+ at the indicated dilution, covered with parafilm, and incubated at 4°C overnight. The next day, cells were washed four times with PBS+ and incubated with the appropriate secondary antibodies conjugated to Alexa488/Alexa594 (Thermo Fisher Scientific) for 1.5-2 hours at room temperature in PBS+. The nuclei were stained using Hoechst 33342 dye (Life Technologies) at 1:2000 dilution in PBS for 5-10 minutes at room temperature. Finally, the cells were washed twice with PBS and mounted in 80% glycerol for imaging.

### Image Acquisition

Images were acquired with Leica SP8 confocal microscope, using LAS-AFX software. Cells were imaged with 63x and 20x objectives. Live cell imaging for Fluorescence Recovery After Photobleaching (FRAP) was performed on a Carl Zeiss780 confocal microscope with an incubation chamber at 37°C and 5% CO_2_. The captured images were processed using Leica Software, Zeiss software, ICY, Fiji/ImageJ, and Adobe CC.

### FRAP analysis

HEK293T cells were grown in 35mm poly-d-lysine coated glass bottom dishes (MatTek) before live cell imaging on a Zeiss780 confocal microscope in an incubation chamber at 37°C and 5% CO_2_. A FRAP region of interest covering a part of an accumulation of GFP-Nup153FG was selected, and imaged for 10 iterations before the FRAP region was bleached for one iteration at the highest intensity of the 488 nm 25mW argon laser focused by a PlanApochromat 63x/1.40 Oil DIC lens. Recovery of fluorescence was monitored at the shortest interval possible (ms range) at 0.5% of the laser intensity used for bleaching. For generation of FRAP curves, the intensity values were normalized to the average intensity of the 10 pre-bleach iteration, and mean±SD were plotted.

### Hexanediol treatment

2 minutes before fixing cells for immunofluorescence, culture medium removed, cells were washed once with 37°C PBS, and new culture medium containing 5% 1,6-hexanediol or 2,5-haxanediol was added.

### 5-Ethynyl Uridine (5-EU) labelling

24 hours prior to fixation of the cells, cells were labelled using medium complemented with 20μM 5-EU (Enzo life sciences). After, the immunofluorescence protocol was followed, but before PBS+ washes, cells were incubated for 30 minutes with 100μl (for 6-well plate) label mix (2mM CuSO4.5H2O, 8μM Sulfo-Cy3 (or other CyColor)-Azide (Lumiprobe), 20mg/ml Ascorbic acid, in PBS), and washed twice with PBS.

### Double thymidine block and FACS sorting for cell cycle analysis

Cells were grown to approximately 40% confluency when synchronised by adding culture medium containing 2mM thymidine for 14 hours. Then cells were washed twice with PBS followed by adding culture medium with 24μM deoxycytidine for 9 hours. The thymidine treatment was repeated for 14 hours before changing to 24μM deoxycytidine again, releasing the second block. Cells were collected by trypsin either immediately or at the indicated time points after release of the block. Trypsin was neutralized by adding culture medium. To determine with FACs the cell cycle phase (G1, S or G2/M) of the synchronized cells, cells were fixed in 70% ethanol overnight at 4°C. DNA content was stained with 50 mg/ml propidium iodide (PI) and 1 mg/ml RNase A for 30 minutes at 37°C in the dark prior to FACS analysis.

### Antibodies and siRNA

The following antibodies and siRNA were used:

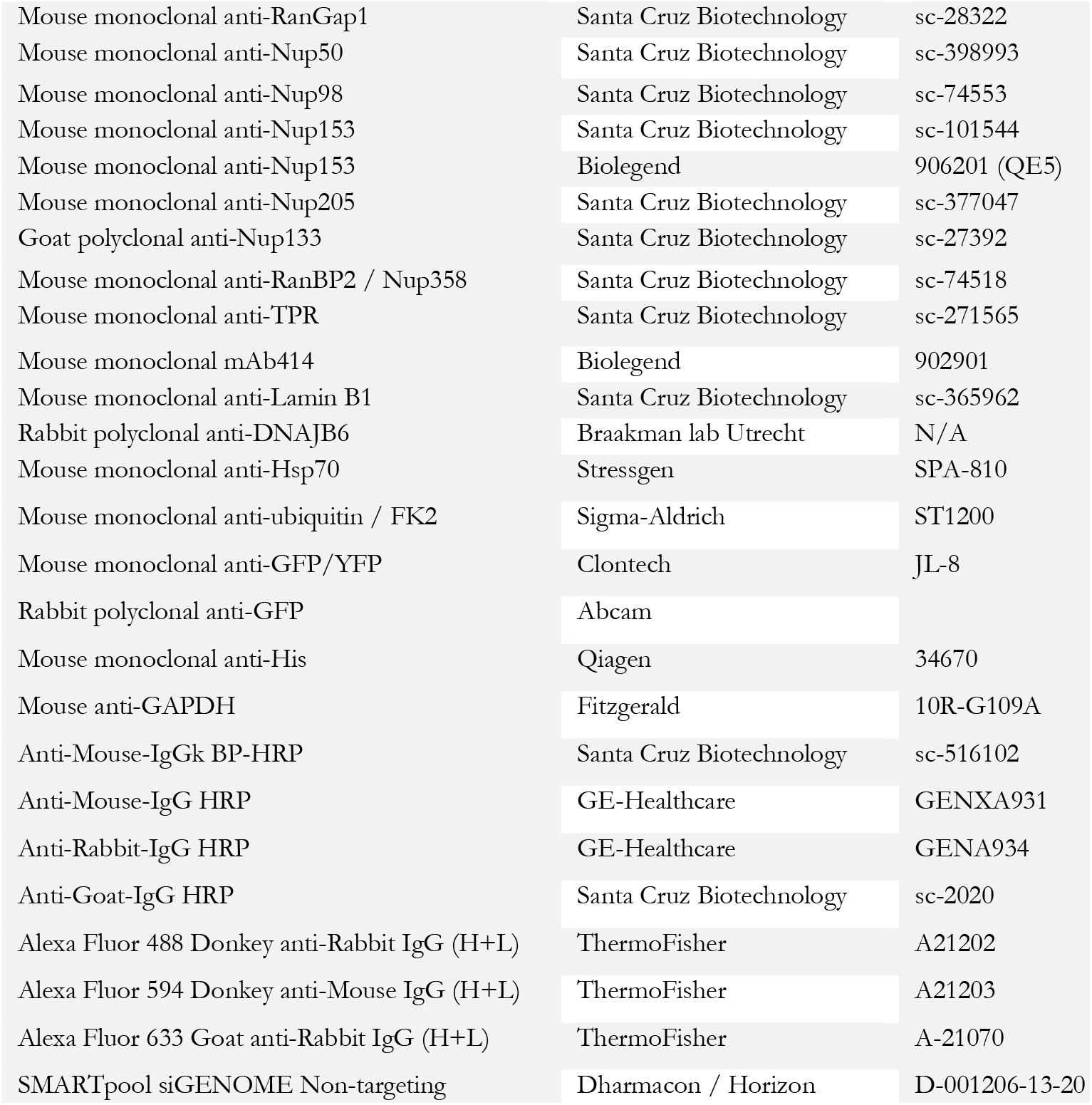
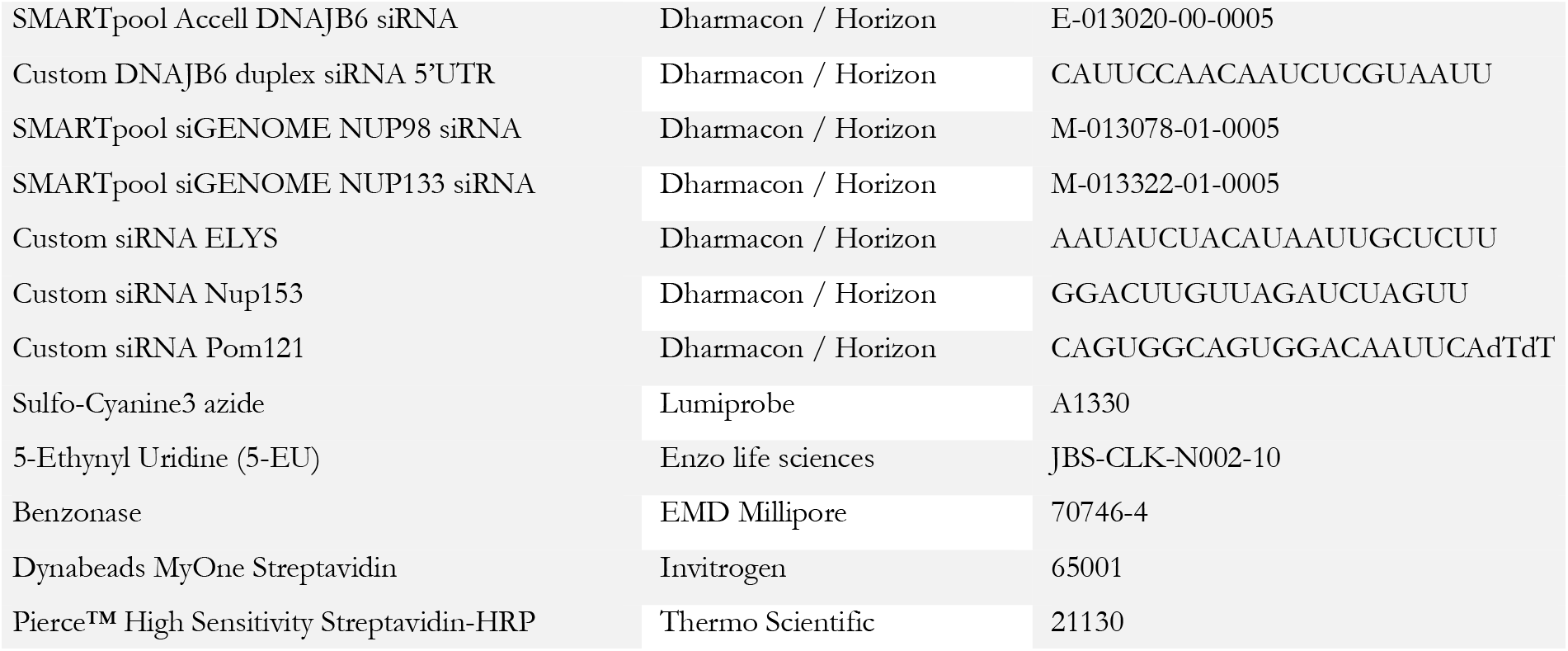

### Transmission electron microscopy

HEK293T DNAJB6 KO cells were fixed in an equal volume of 2% PFA and 0.2% GA in 0.1 M sodium cacodylate buffer (pH 7.4) in medium and incubated for 10 min. Cells were then incubated with fresh pure fixative (2% PFA and 0.2% GA) for 30 min and rinsed twice with 0.1 M sodium cacodylate. All steps were performed at RT. Epon EM embedding was performed as described previously^56^.

### Immuno-electron microscopy

HEK293T cells with CRISPRi knockdown of Torsin A and B, with or without overexpression of GFP-DNAJB6b, were fixed by adding an equal volume of culture medium of freshly prepared double strength fixative (4% formaldehyde, 0,4% glutaraldehyde in 0.1 M phosphate buffer, pH 7.4) for 5 minutes at room temperature (RT). This mixture was then replaced by one volume of single strength fixative (2% formaldehyde, 0.2% glutaraldehyde in 0.1M phosphate buffer, pH 7.4) before incubating the cells for additional 3 hours at RT. Cell embedding, ultrathin cryosectioning (50-60 nm) and immunogold labelling were performed as previously described^36^. Sections were immuno-labelled using either a rabbit anti-GFP (Abcam) or an anti-DNAJB6 antibodies, which were subsequently detected with protein A-gold (Cell Microscopy Center, UMC Utrecht, the Netherlands). For electron tomography, additional fiducial markers (15nm protein A colloidal gold, Cell Microscopy Center, UMC Utrecht, the Netherlands) were layered on top of the thick cryosections after the immunogold labelling. For 2D imaging, cell sections were analysed using an 80 kV transmission electron microscope CM100bio TEM (FEI, Eindhoven, Netherlands).

### Electron tomography: 3D reconstruction and modelling

Three-dimensional models of GFP-DNAJB6b-positive and endogenous DNAJB6-postitive herniations were obtained by electron tomography of 300- to 400-nm thick cryosections. The areas of interest were selected based on the immuno-gold detection of GFP-DNAJB6b, using a conventional 80 KV electron microscope, prior recording of the double-tilt series with high voltage (200KV) electron microscope. The tilt series images, recorded using a Tecnai T20 (FEI), were aligned using at least 20 fiducial gold particles, using the IMOD program package (University of Colorado, USA). The IMOD software was also used to create double tilt tomograms by combining two R-weighted back projection and/or SIRT tomograms. Filtering options in the IMOD package (median algorithm) were used to ‘smooth’ the tomograms. The tomograms had a final lateral resolution of approximately 4 nm based on the Crowther criterion. Double tilt tomograms were analysed and modelled also using the IMOD software. Features of interest were contoured manually in serial optical slices extracted from the tomogram, and used to create surface-rendered models.

### Western blot analysis

Indicated sample amounts and 5μl PageRuler Prestained Protein Ladder (Thermo Scientific) were loaded on 10% or 12% SDS-PAGE gels. SDS-PAGE was performed using the BioRad Mini-PROTEAN 3 system. After SDS-PAGE, proteins were transferred to nitrocellulose membranes by semiwet electrotransfer with the Bio-Rad Trans-Blot Turbo transfer system. To prevent non-specific protein binding, membranes were incubated in 5% (w/v) non-fat dried milk in PBS-Tween (PBS-T) for one hour at room temperature. Membranes were washed in PBS-T and incubated at 4°C overnight with indicated antibodies. Membranes were washed three times 10 minutes and incubated for at least one hour in HRP-conjugated secondary antibodies (GE Healthcare) at 1:8000 dilution in PBS-T. After incubation, membranes were washed three times 10 minutes in PBS-T and enhanced chemiluminescence detection was performed using the ECL Western Blotting substrate kit (Thermo Scientific). Quantifications were performed using Bio-Rad imaging software and graphs were generated using GraphPad prism. Statistical significance was analysed using an independent Student’s t-test, p<0.05 was considered statistically significant. Values are expressed as the mean ± SD.

### Protein extraction for Filter Trap Assay

48 hours after transfection cells were washed once in cold phosphate-buffered saline (PBS) and scraped in 200μl lysis buffer (25 mM HEPES, 100 mM NaCl, 1 mM MgCl_2_, 1% NP40 (Igepal CA-630, Sigma), EDTA free complete protease inhibitors cocktail (Roche), Benzonase (EMD Millipore) (~90 units/ml)), and left on ice for minimum of 30 minutes with intermittent vortexing until chromatin was dissolved. Protein concentrations were determined using DC protein assay (Bio-Rad). Concentrations were equalized and diluted in FTA buffer (10 mM Tris-HCl pH 8.0, 150 mM NaCl and 50 mM dithiothreitol, 0.5% SDS), boiled for 5 minutes and prepared in three 5-fold serial dilutions into a final of 1x, 5x and 25x diluted samples for filter trap assay (FTA). FTA samples were loaded onto a 0.2 μm pore size cellulose acetate membrane prewashed with 0.1% SDS-containing FTA buffer. Membranes were washed two times with 0.1% SDS-containing FTA buffer, blocked with 5% non-fat milk, and blotted with anti-GFP/YFP (Clontech). After HRP-conjugated secondary antibody incubation, visualization was performed using enhanced chemiluminescence and ChemiDoc Imaging System (Bio-Rad). Bands were measured (ImageLab 5.2.1.) and the measurement of the three dilutions was averaged. Values normalized to control were plotted in a graph.

### Filter trap assay purified proteins

To assay the effect of crowding agents in the aggregation/phase transition of FG Nups, purified proteins were diluted at the indicated concentrations in the assay buffer (50mM TrisHCl 150mM NaCl, pH 8.0), containing varying concentrations of the corresponding crowding agent: 0/5/10/305515% w/v of serine (L-Serine, 99%, ACROS Organics™, AC132660250), 0/5/10/15% w/v of Polyethylene glycol 3350 (PEG, Sigma-Aldrich, P4338-2KG), 0/5/10/15% w/v of Ficoll (Sigma-Aldrich, F2637-25G). To assay the effect of the chaperone DNAJB6b in the aggregation/phase transition of FG-Nup fragments, purified FG-fragments were mixed with purified DNAJB6b, or the corresponding control protein (BSA, MBP-mCherry, DNAJA1), in the assay buffer (50mM TrisHCl 150mM NaCl, pH 8.0) containing 10% PEG. Aggregation assays were performed with a protein concentration of 3μM in a final volume of 20μl, at 25°C for either 1 hour or 3 hours. While for hNup153FG and yNup100FG 1 hour was sufficient to form SDS-insoluble aggregates in assay buffer, yNup116FG required 3 hours to form aggregates. For yNup145N the buffer composition was adjusted to 50mM NaCl, 100mM sodium phosphate, pH 6 containing 10% PEG3350 (w/v) to allow for the formation of SDS-insoluble aggregates (at 25°C for 1 hour).

For filter trap assay, 3 trans-blot papers (Bio-Dot SF Filter Paper, Bio-Rad, #1620161) and one cellulose acetate membrane (Cellulose Acetate Membrane Filters, 0.2 Micron, Sterlitech, CA023001) were soaked in either 150mM NaCl, 50mM TrisHCl pH 8.0 (hNup153FG, yNup100FG, yNup116FG) or 50mM NaCl, 100mM sodium phosphate pH 8.0 (yNup145N FG) buffer, placed on the Bio-Dot apparatus (Bio-Rad, #1703938), and washed four times with sample buffer (150mM NaCl, 50mM TrisHCl, pH 8.0/50 mM NaCl, 100 mM sodium phosphate pH 8.0), applying constant vacuum. Samples were added to 180μL of sample buffer supplemented with 0.5% SDS and mixed by vortex before loading in the Bio-Dot apparatus (Bio-Rad). After loading samples and applying constant vacuum, slots were washed twice with sample buffer (50mM TrisHCl 150mM NaCl, pH 8.0/50mM NaCl, 100mM sodium phosphate pH 8.0) and twice with wash buffer (150mM NaCl 50mM TrisHCl pH 8.0/50mM NaCl, 100mM sodium phosphate pH 8.0 supplemented with 0.1% SDS). After blocking for 1 hour in 2.5% BSA (Albumine bovine serum, Acros Organics 268131000, #10450141) in PBS with 0.1% Tween20 (MP Biomedicals™, MP1TWEEN201), the membrane was incubated with mouse primary antibody α-His (1:5000, Monoclonal mouse Tetra-His antibody, Qiagen, #34670) in 2.5% BSA in PBS-T overnight at 4°C, washed three times with PBS-T, incubated with anti-mouse secondary antibody m-IgGk BP-HRP (1:5000, Santa Cruz Biotechnology, sc-516102) in 2.5% BSA in PBS-T for 1 hour at room temperature, and washed three times with PBS-T. Chemiluminescence was detected using ECL substrate (GE Healthcare, RPN2232) according to manufacturer’s instructions on a ChemiDoc Touch Imaging System (Bio-Rad).

### Protein purification

The expression of FG-Nup fragments, DNAJB6b, DNAJA1, and MBP-mCherry was performed in 200ml cell cultures of *Escherichia coli* cells (OD600=0.5-0.8, grown at 37°C) by adding 0.5mM IPTG (Isopropyl-β-D-thiogalactoside, #10724815001) for 5 hours at 20°C. Cells were harvested by centrifugation (4500g, 15 minutes, 4°C) and pellets were stored at −80°C. For the purification of FG-fragments and the chaperones DNAJB6 and DNAJA1, the cell pellets were resuspended in 10ml lysis buffer (100mM Tris-HCl 2M Guanidine-HCl buffer, pH 8.0) (Thermo Scientific™,# 24110), supplemented with protease inhibitor cocktail (cOmplete™ ULTRA Tablets, Mini, EDTA-free, EASYpack Protease Inhibitor Cocktail, #5892791001), 1mM PMSF, (Phenylmethylsulfonyl fluoride, Serva, #32395.02), and 5mM DTT (DL-1,4-Dithiothreitol, 99%, ACROS Organics™, #10215550), and disrupted with glass beads (disruptor beads 0.1mm, Scientific Industries, SI-BG01) in the Fastprep. Lysed cells were clarified by centrifugation for 5 minutes at 13.000rpm and the supernatant was centrifuged again for 15 minutes at 13.000rpm. The supernatant was equilibrated in lysis buffer and incubated with Ni-sepharose beads (Ni Sepharose® 6Fast Flow, Cytiva, 17-5318-03) at 4°C for 1 hour. Beads were washed three times with 10ml of lysis buffer (100mM Tris-HCl 2M Guanidine-HCl, pH 8.0) supplemented with 50mM Imidazole (PUFFERAN® ≥99 %, CarlRoth, X998.4) and 2.5mM DTT, using Poly-Prep columns (Poly-Prep® Chromatography Columns, Bio-Rad, #7311550). Proteins were eluted with 500mM Imidazole (100mM Tris-HCl 2M Guanidine-HCl, pH 8.0). A 5μl fraction of the purified proteins was checked on SDS-PAGE electrophoresis (Fig. S6). Cysteine residues of FG-Nup fragments were reduced with 10mM DTT for 30 minutes at 50°C and blocked with 15mM iodoacetimide alkylating reagent (Sigma Aldrich, #I1149-25G) for 30 minutes at room temperature to prevent the formation of disulfide bonds. Proteins were concentrated using Vivaspin Protein Concentrator spin columns (Vivaspin® 2, 10/30 kDa MWCO Polyethersulfone, Cytiva, 28-9322-47, 28-9322-48), according to manufacturer’s instructions. Expression and purification of MBP-mCherry was performed essentially as described before, but using 50mM Tris-HCl 150mM NaCl, pH 8.0 buffer for cell lysis and protein purification. Measurement of final protein concentration was performed using BCA protein assay (Pierce™ BCA Protein Assay Kit, Thermo Scientific™, #23227), according to manufacturer’s instructions, using BSA (Pierce™ Bovine Serum Albumin Standard Ampules, 2mg/mL, # 23209) as a standard.

## Data availability

The authors declare that all data supporting the findings of this study are available within the article and its supporting information files or from the corresponding author upon reasonable request.

---

**Supplementary Figure 1.**
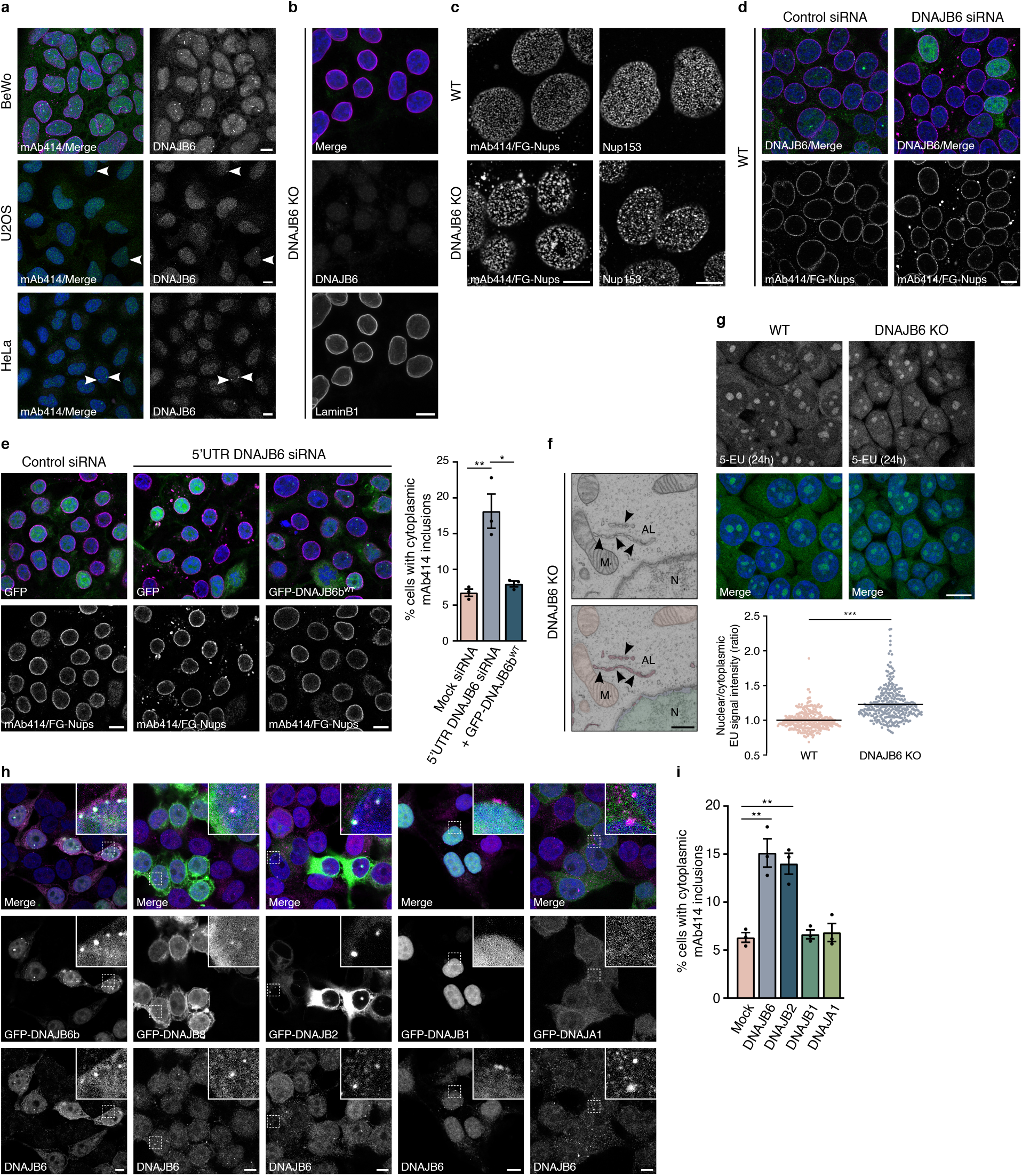
Disruption of DNAJB6 expression leads to accumulation of nuclear pore complexes (NPCs) in the cytoplasm. **A.** Different cell lines stained for DNAJB6 (green) and FG-Nups (mAb414, magenta). BeWo cells show high amount of DNAJB6b foci. U2OS and HeLa cells show almost no DNAJB6 foci. **B.** Z-projection of 20 consecutive slices of 0.5μm of HEK293T DNAJB6 KO cells stained for DNAJB6 (green) and LaminB1 (red). **C.** Representative high-resolution confocal images of mAb414 or Nup153-stained nuclear pores (focused on the top of the nucleus) of HEK293T and HEK293T DNAJB6 KO cells show aberrant NPC distribution in the NE of HEK293T DNAJB6 KO cells. **D.** siRNA-mediated knockdown of DNAJB6 (green) (72h) induces cytoplasmic accumulations of NPCs, or annulate lammelae (AL), stained by mAb414 (magenta). **E.** HEK293T cells transfected with siRNA against the 5’UTR of DNAJB6 show AL and by simultaneous expression of GFP-DNAJB6b^WT^, containing only the cDNA, AL formation can be prevented. With a quantification of AL formation after knockdown of DNAJB6.*p≤0.05, **p≤0.01, n=3 independent samples, unpaired t-test, mean±SEM. **F.** AL visualised by transmission electron microscopy and pseudo-colored in bright pink, can be distinguished in cytoplasmic strings. Arrows point at individually identifiable NPCs. Nucleus indicated with N, mitochondrion indicated with M. Scale bar represents 500nm. **G.** Representative images of HEK293T and HEK293T DNAJB6 KO cells cultured in medium containing the uridine analog 5-ethynyluridine (5-EU) for 24 hours. 5-EU incorporation into newly transcribed RNA was detected by Cy3-azide (green). The ratio of nuclear to cytoplasmic signal intensities of RNA per cell, each individual point represents a single cell. ***p<0.001, unpaired t-test. >300 cells were counted over 3 independent experiments. **H.** Colocalization of GFP-tagged DNAJB6b, DNAJB8, and DNAJB2 with endogenous DNAJB6. DNAJB1 and DNAJA1 do not colocalize with DNAJB6 foci at the NE. **I.** AL formation under siRNA-mediated knockdown of DNAJB6, DNAJB2, DNAJB1, and DNAJA1. Knockdown of DNAJB6 and DNAJB2 leads to significant increase in AL formation. *p≤0.05, **p≤0.01, unpaired t-test; n=3 independent samples with total 4000-5000 cells counted, mean±SEM. Scale bars on all fluorescent images represent 10μm.

**Supplementary Figure 2.**
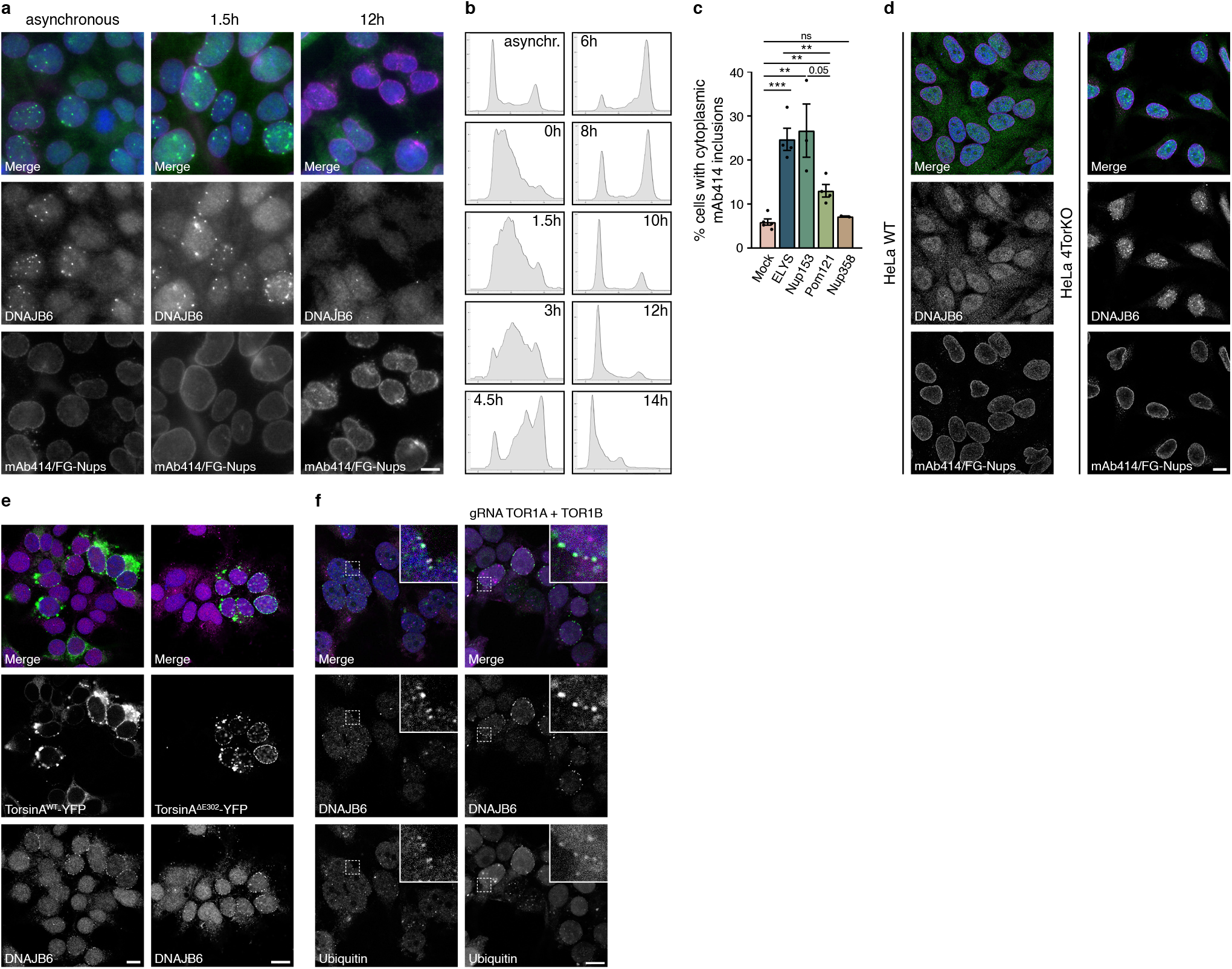
DNAJB6 foci formation is cell cycle dependent and is related to interphase NPC assembly. **A.** DNAJB6 foci in HEK293T cells in asynchronized cells, and cells 1.5h and 12h after release from a double thymidine cell cycle synchronization. **B.** Cell cycle progression was analysed by fluorescence-activated cell sorting (FACS). Cells were stained with propidium iodide to analyze DNA content. **C.** Quantification of percentage of cells with cytoplasmic NPC accumulations (AL) stained with mAb414 under siRNA knockdown of ELYS, Nup153, Pom121, or Nup358. **p≤0.01, ***p≤0.001; n≥3 independent samples, mean±SEM. **D.** HeLa WT and HeLa 4TorKO cells stained for DNAJB6 (green) and FG-Nups with mAb414 (magenta). DNAJB6 only localizes to few foci in HeLa WT cells, but HeLa 4TorKO have many DNAJB6 foci. **E.** Overexpression of either TorsinA wildtype or ΔE302 (green) leads to an increase in foci at the nuclear envelope, containing DNAJB6 (magenta). **F.** Ubiquitin (magenta) colocalizes with DNAJB6 (green) at foci at the nuclear envelope and are increased upon torsin knockdown (gRNA vector containing mTurquoise2). Scale bars on all fluorescent images represent 10μm.

**Supplementary Figure 3.**
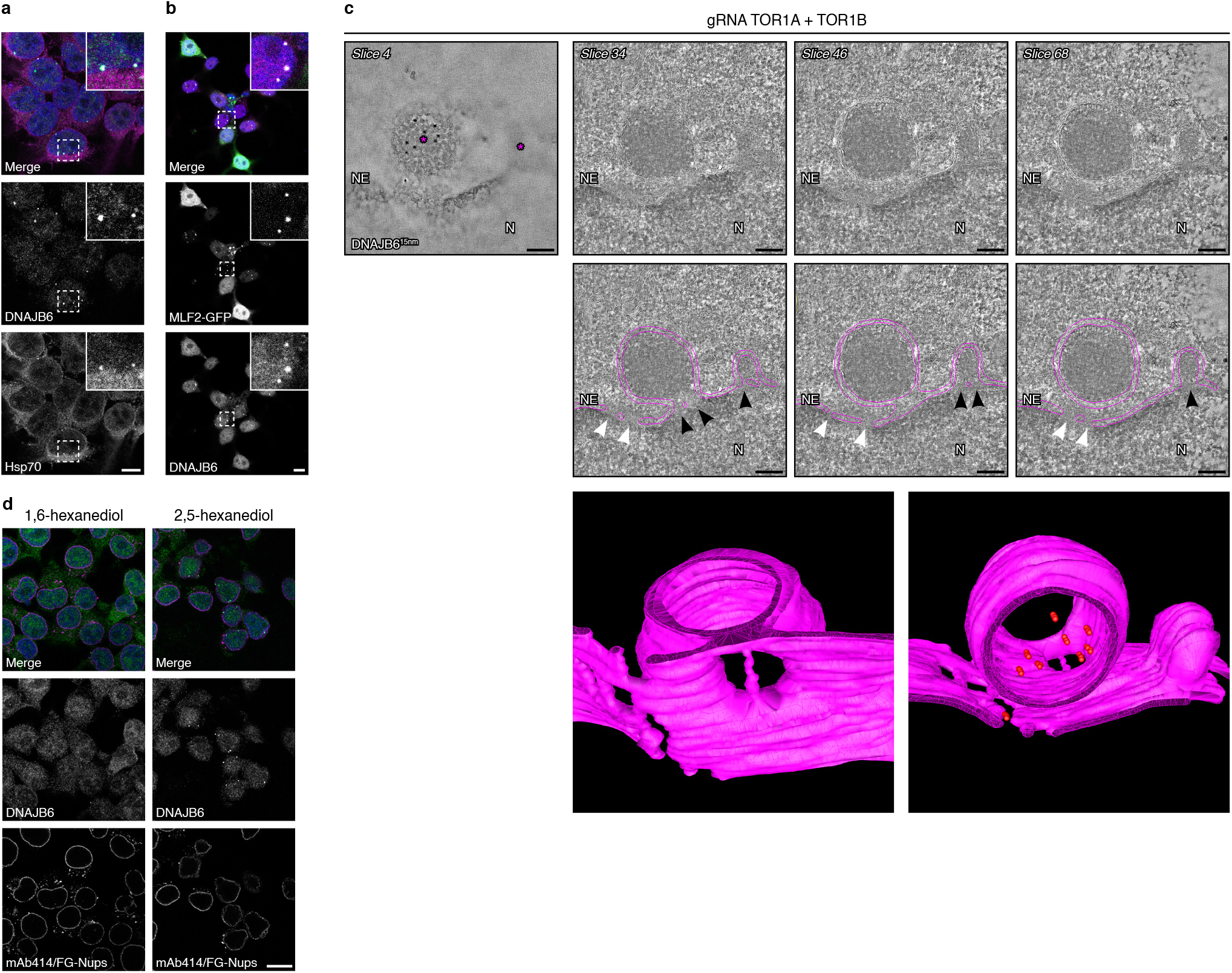
DNAJB6 localizes to herniations at the NE and is dependent on intact NPCs. **A.** HSP70 (magenta) colocalizes with DNAJB6 (green) in foci at the NE. **B.** MLF2-GFP colocalizes with DNAJB6 (magenta) in foci at the NE. **C.** For immunoelectron tomography, surface labelling was used to select areas of interest prior to recording tilt series. Tomographic slice (Z slice 4) revealing the specific immunogold (15 nm) labelling on the surface of the cryosection. Asterisk indicating herniations. Indicated tomographic slices extracted from the tomogram showing herniations at the NE. In the bottom images, nuclear membranes are traced (pink). More electron dense areas can be observed at the neck of the herniations where the NE bends into the herniation, indicated by black arrow heads. NPCs are indicated with white arrow heads. Nucleus is indicated with N and nuclear envelope with NE. Scalebars represent 232nm. Lower images are from the three-dimensional model of DNAJB6-positive herniations, obtained by electron tomography of 400 to 450 nm thick cryosections. The NE is represented in pink. Left is a view from the nuclear side showing the openings of the NPCs and of the herniations. Right shows the herniations with the localization of the DNAJB6 gold labelling in red. **D.** Cells treated with 5% 1,6-hexanediol (2 minutes) lose almost all DNAJB6 foci (green), while cells treated with 2,5-hexanediol have only a slight reduction in foci. Scale bars on all fluorescent images represent 10μm.

**Supplementary Figure 4.**
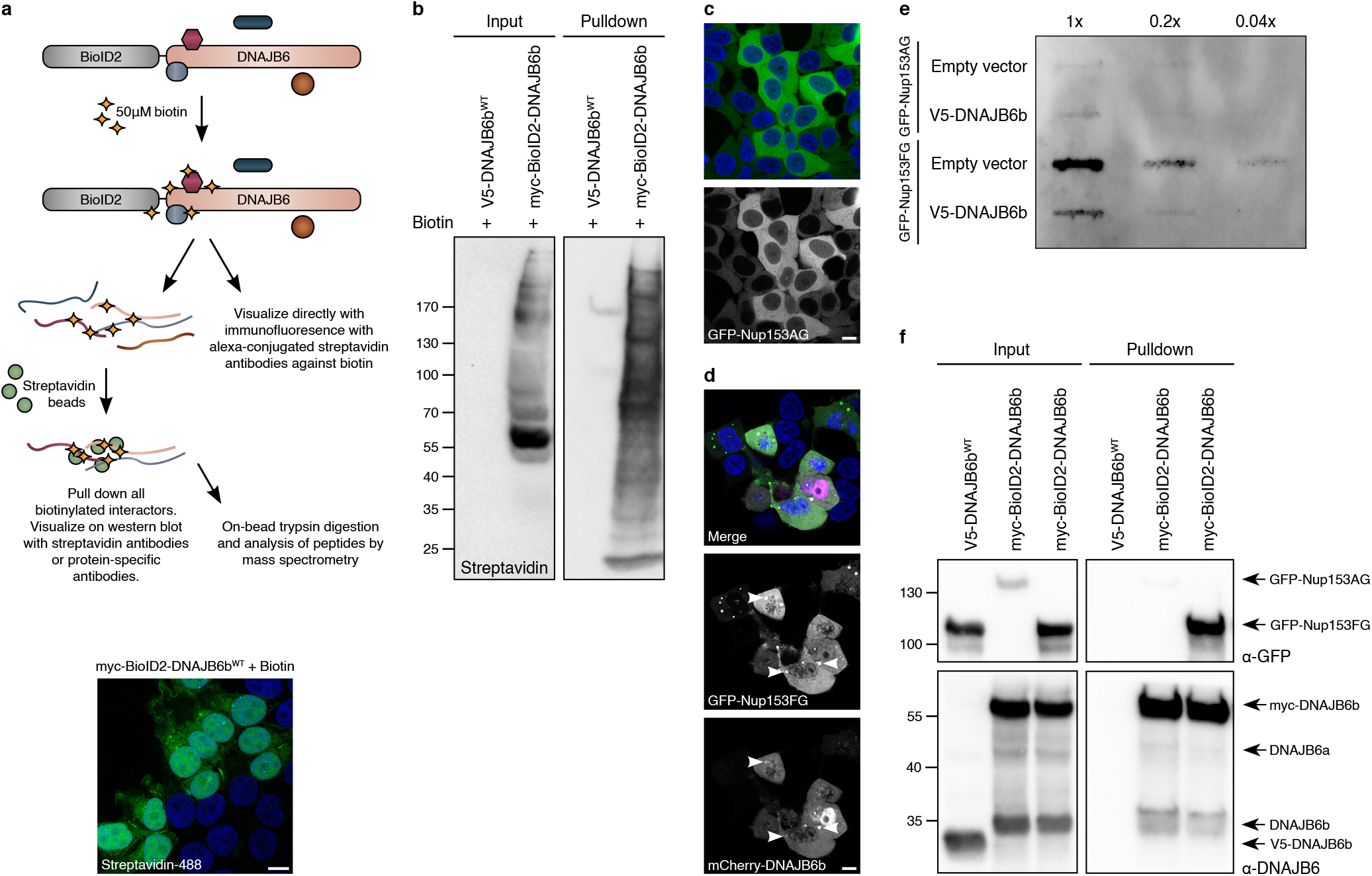
The interaction of DNAJB6 with hNup153FG depends on the FG-motifs in hNup153FG. **A.** Schematic overview of the BioID2-DNAJB6b fusion protein and how samples were obtained. Fluorescent image shows HEK293T cells expressing BioID2-DNAJB6b, treated for 24h with 50μM biotin and stained with Alexa Fluor 488-conjugated streptavidin, biotinylated substrates are visualized in foci at the nuclear envelope. **B.** Streptavidin blot belonging Fig. 4A where HEK293T cells expressing V5-DNAJB6b or BioID2-DNAJB6b are treated for 24h with 50μM biotin. HRP-conjugated streptavidin binds to all biotinylated close proximity partners of DNAJB6b that are pulled down by biotin affinity immunoprecipitation with streptavidin beads and run on SDS-PAGE. **C.** GFP-Nup153AG overexpression in HEK293T cells is homogeneously distributed in the cytoplasm. Scale bars on all fluorescent images represent 10μm. **D.** HEK293T cells overexpressing GFP-Nup153FG and mCherry-DNAJB6 show colocalization in spherical accumulations (arrowheads). **E.** Filter-trap assay of HEK293T cells with GFP-Nup153AG or GFP-Nup153FG overexpression. Three serial 5-fold dilutions were loaded onto cellulose-acetate membranes and probed with anti-GFP antibodies to detect aggregation of the GFP-Nup153FG or GFP-Nup153AG fragment without or with co-overexpression of V5-DNAJB6b. **F.** HEK293T cells overexpressing V5-DNAJB6b or BioID2-DNAJB6b, either with GFP-Nup153FG or GFP-Nup153AG, are treated for 24h with 50μM biotin. Biotinylated close proximity partners of DNAJB6b are pulled down by biotin affinity immunoprecipitation with streptavidin beads, run on SDS-PAGE. GFP and DNAJB6 are detected with specific antibodies. Despite the lower levels of Nup153AG, the relative intensities in the pulldown indicate that binding of DNAJB6 is dependent on the FG-motifs (lane 2 and 3 of the pulldown blot).

**Supplementary Figure 5.**
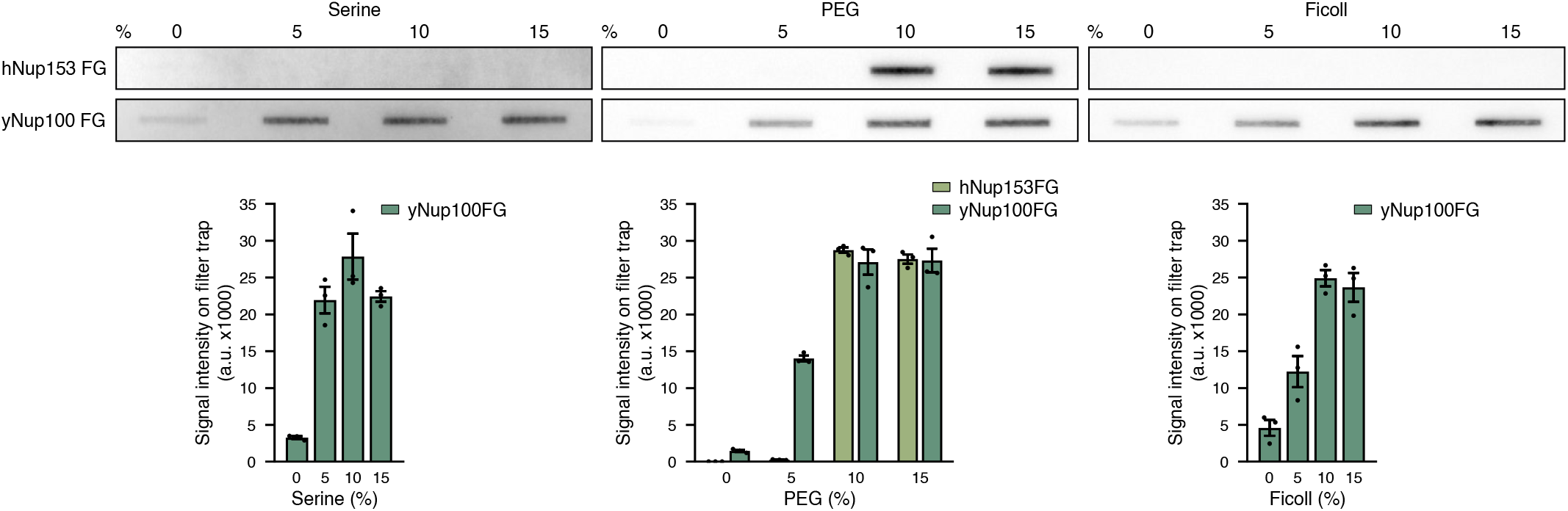
The effect of crowding agents on the aggregation of human Nup153FG and yeast Nup100FG. Top: 3μM of purified FG-Nup fragments were incubated in assay buffer (150mM NaCl, 50mM TrisHCl, pH8) containing the indicated concentration (w/v) of the crowding agents Serine, PEG3350, or Ficoll, at 25°C during 1 hour, and loaded on a filter trap. Bottom: Quantification of Nup band intensity in the described conditions; hNup153FG did not form any quantifiable bands on filter trap with the addition of Serine or Ficoll. n=3 independent samples; mean±SEM.

**Supplementary Figure 6.**
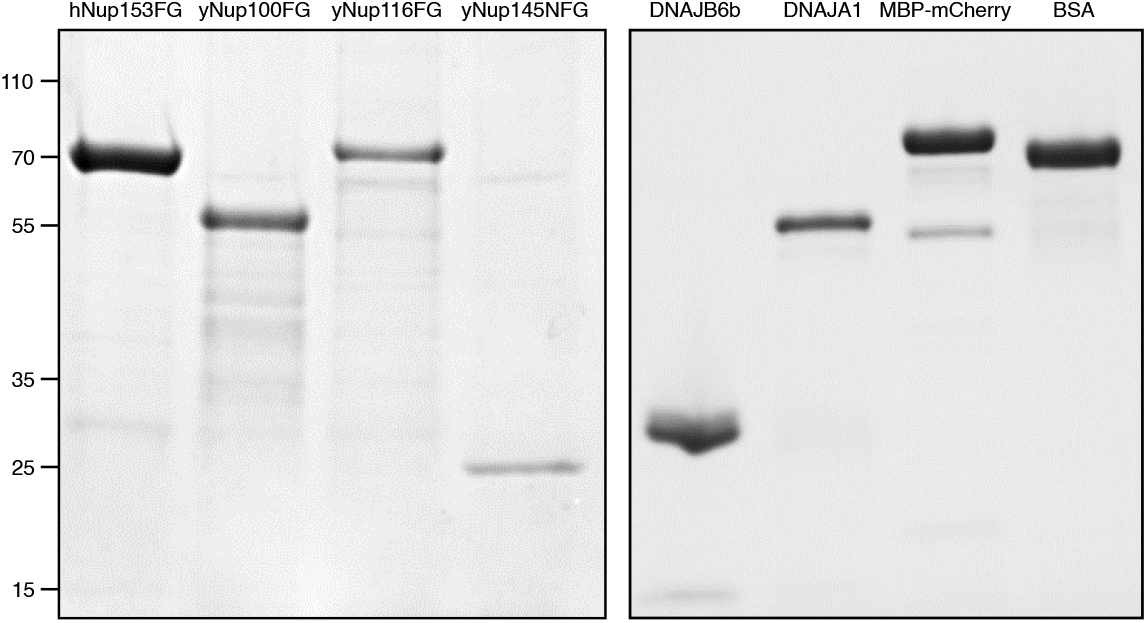
Coomassie Brilliant Blue stained SDS-PAGE of indicated purified protein samples.

